# Seasonal and annual fluctuations of deer populations estimated by a Bayesian state–space model

**DOI:** 10.1101/844688

**Authors:** Inoue Mizuki, Hiroki Itô, Michimasa Yamasaki, Shigeru Fukumoto, Yuuki Okamoto, Masaya Katsuki, Keitaro Fukushima, Masaru Sakai, Shota Sakaguchi, Daisuke Fujiki, Hikaru Nakagawa, Masae Ishihara, Atsushi Takayanagi

## Abstract

Deer overabundance is a contributing factor in the degradation of plant communities and ecosystems worldwide. The management and conservation of the deer-affected ecosystems requires us to urgently grasp deer population trends and to identify the factors that affect them. In this study, we developed a Bayesian state–space model to estimate the population dynamics of sika deer (*Cervus nippon*) in a cool-temperate forest in Japan, where wolves (*Canis lupus hodophilax*) are extinct. The model was based on field data collected from block count surveys, road count surveys by vehicles, mortality surveys during the winter, and nuisance control for 12 years (2007–2018). We clarified the seasonal and annual fluctuation of the deer population. We found two peaks of deer abundance (2007 and 2010) over 12 years. In 2011 the estimated deer abundance decreased drastically and has remained at a low level then. The deer population increased from spring to autumn and decreased from autumn to winter in most years. The seasonal fluctuation we detected could reflect the seasonal migration pattern of deer and the population recruitment through fawn births in early summer. In our model, snowfall accumulation, which can be a lethal factor for deer, may have slightly affected their mortality during the winter. Although we could not detect a direct effect of snow on population dynamics, snowfall decrease due to global warming may decelerate the winter migration of deer; subsequently, deer staying on-site may intensively forage evergreen perennial plants during the winter season. The nuisance control affected population dynamics. Even in wildlife protection areas and national parks where hunting is regulated, nuisance control could be effective in buffering the effect of deer browsing on forest ecosystems.

## Introduction

In the past few decades, deer have become increasingly abundant worldwide [1, 2]; this population increases has contributed to the degradation of plant communities and ecosystems [3 – 6]. In general, the population dynamics of animals are affected by birth, mortality, and migration rates. Large ungulates are able to breed under low food availability [7], therefore, the birth rate of deer would not largely decrease even in a degraded forest; however, the density-dependent decline in the birth rate of deer occurs at a later period of the outbreak stage [8]. Furthermore, the survival rate of adult deer was high even in a poor nutritional environment [9]. Thus, deer is a species that can live in high densities and low-nutrient environments. If predators (e.g. wolves) are absent, hunting is one options to control deer populations under these conditions [10, 11].

In snow-covered area and, in particular, during heavy snowfall, the survival rate of sika deer (*Cervus nippon*) decreases [9, 12]. Kawase et al. (2014) [13] projected that winter precipitation including snowfall would decrease in broad regions of Japan due to the ongoing climate change. This climate change may mitigate the mortality of deer and cause further increases in deer populations in the future. Therefore, it is indispensable to estimate the effect of snow on the dynamics of deer populations. While some of the effects of global warming on population dynamics of ungulates have already been reported [14–16], models constructed in recent studies to describe deer population dynamics have not yet explicitly considered the effects of snow.

From the viewpoint of plant communities and ecosystems, it is important to clarify not only annual trends but also seasonal trends in the deer population density. Plant fitness could be affected differently depending on whether deer browse on them before or after they have reproduced sexually. The timing of browsing could also affect the fitness of pollinators such as bumblebees. Therefore, in order to assess the effects of deer browsing on ecosystem levels it is important to, at least, estimate the seasonal deer abundance. However, in many areas, the annual census of ungulates is held during a season that, although offers good visibility to track ungulates, is not suitable for plant growth [10, 15, 17].

In recent years, generalized linear models [18], generalized additive mixed models (GAMM, [19]), density surface models [20], and Bayesian state-space models [10, 16] were used to estimate deer abundance based on field data. Among these models, the Bayesian state-space model can be a powerful tool for estimating deer population dynamics because it can easily handle time series data with temporal autocorrelation and can explicitly distinguish errors following measurement of data with uncertainty about population dynamics [10, 21]. However, there are still limited applications of this model when it comes to the effect of snow and seasonal fluctuations on deer population dynamics.

In this study, we estimated deer population dynamics in a cool-temperate forest in Japan using a Bayesian state-space model. The model was based on data collected from block count surveys, road count surveys by vehicles, mortality surveys during the winter, and nuisance control over 12 years. The seasonal and annual fluctuation of the sika deer population and the effects of snowfall and nuisance control on population dynamics are discussed based on the results we obtained by the model and the parameters estimated in the model, respectively.

## Materials and Methods

### Study site

The study site was located at the Ashiu Forest Research Station, Field Science Education and Research Center, Kyoto University, Japan (35°20’N, 135°45’E; 355 – 959 m a.s.l., 4,186 ha) and the surrounding area (4,612 ha in total, Fig 1). The mean annual temperature and precipitation in this area are 13.1 C and 2,333 mm, respectively [22]. The maximum snowfall during each winter at 356 m elevation was 31.0 – 141.7 cm between 2007 and 2018 (Table 1). The forest is usually closed from January to early April because the roads in the forest must be blocked with snow. This forest is located in the transition part between the temperate deciduous forest zone and the warm temperate forest zone. This area is well known for being highly diverse in plant species and existing phylogeographically important populations of some species in the forest [22]. Though the forest is one of the wildlife protection areas in Japan, forest vegetation has been steadily degraded by the browsing of *C. nippon* [23, 24]; thus, nuisance control started in 2008 using guns, traps, and cages (Table 2). The last known Japanese wolf (*Canis lupus hodophilax*) was caught in the Nara prefecture in 1905 and there have been no sightings of it in Japan since. Thus, we considered that potent predators of deer such as wolves had been extinct all over Japan, including in our study site.

**Fig 1.**
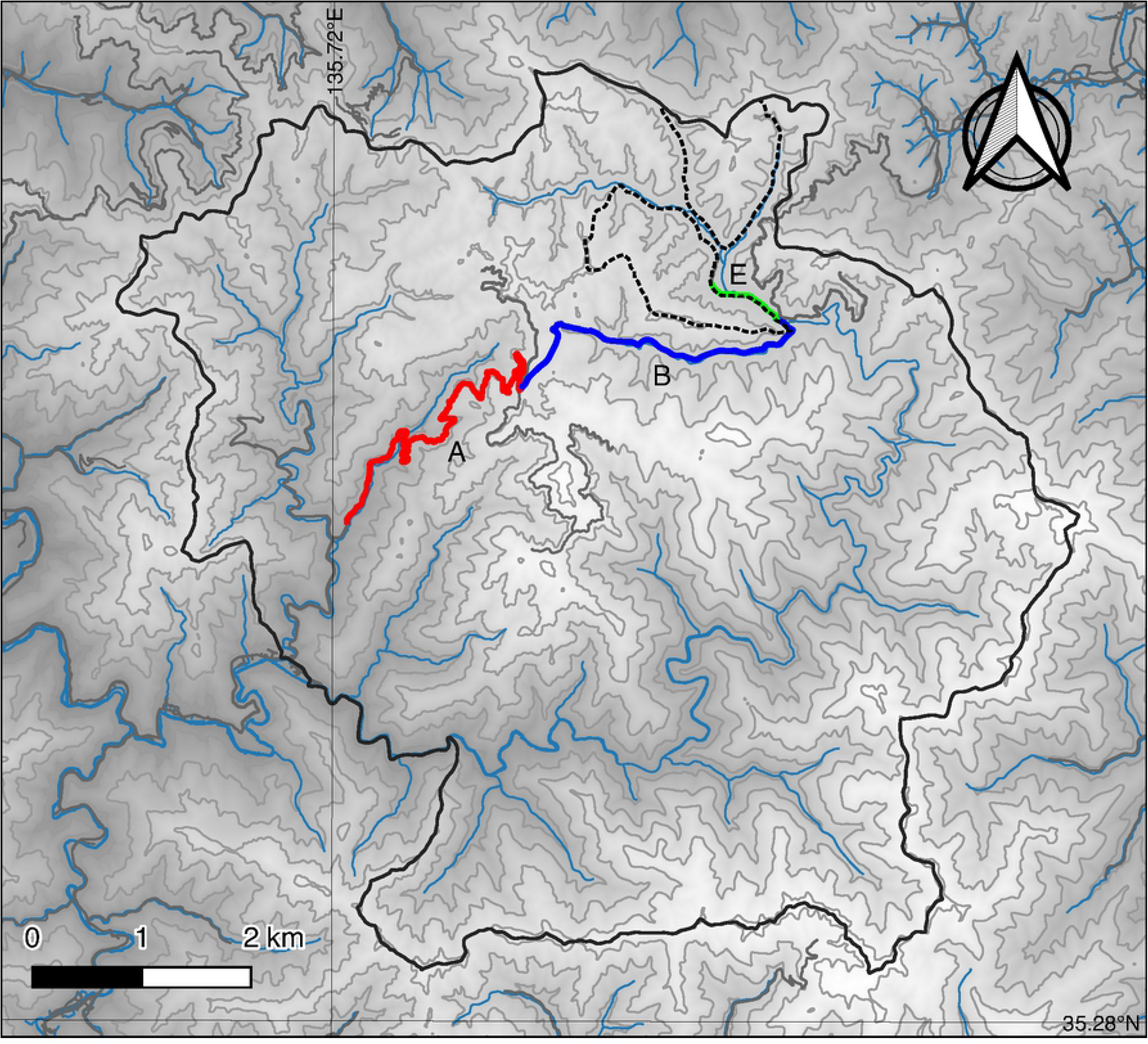
Topographic map of Ashiu Forest Station and the surrounding area. Red, blue, and green lines denote the location of the selected route sectors, A, B, and E, respectively. The parts surrounded by solid lines denote the area of Ashiu Forest Station and the area surrounding. The parts surrounded by broken lines denote survey area by block count.

**Table 1.** The numbers of deer carcasses in winter and winter climate.

| Year | Carcasses<sup>a</sup> | Distance (km)<sup>b</sup> | SD50<sup>c</sup> (days) | MaxS<sup>d</sup> (cm) |
| --- | --- | --- | --- | --- |
| Nov 2007 - Mar 2008 | 30 | 78.5 | 52 | 132 |
| Nov 2008 - Mar 2009 | 0 | 85.0 | 40 | 130 |
| Nov 2009 - Mar 2010 | 0 | 98.5 | 0 | 36.7 |
| Nov 2010 - Mar 2011 | 25 | 88.5 | 76 | 141.7 |
| Nov 2011 - Mar 2012 | 4 | 83.5 | 87 | 138.3 |
| Nov 2012 - Mar 2013 | 7 | 101.5 | 21 | 90.7 |
| Nov 2013 - Mar 2014 | 6 | 83.5 | 70 | 114 |
| Nov 2014 - Mar 2015 | 38 | 100.0 | 71 | 113 |
| Nov 2015 - Mar 2016 | 2 | 92.5 | 0 | 31 |
| Nov 2016 - Mar 2017 | 30 | 86.5 | 61 | 135 |
| Nov 2017 - Mar 2018 | 0 | 93.0 | 7 | 87 |

<sup>a</sup>Number of deer carcasses
<sup>b</sup>Distance of the total survey area during spring thaw
<sup>c</sup>Snow cover >50 cm duration
<sup>d</sup>Maximum snow depth at 356 m elevation at the Ashiu Forest Station.

**Table 2.**
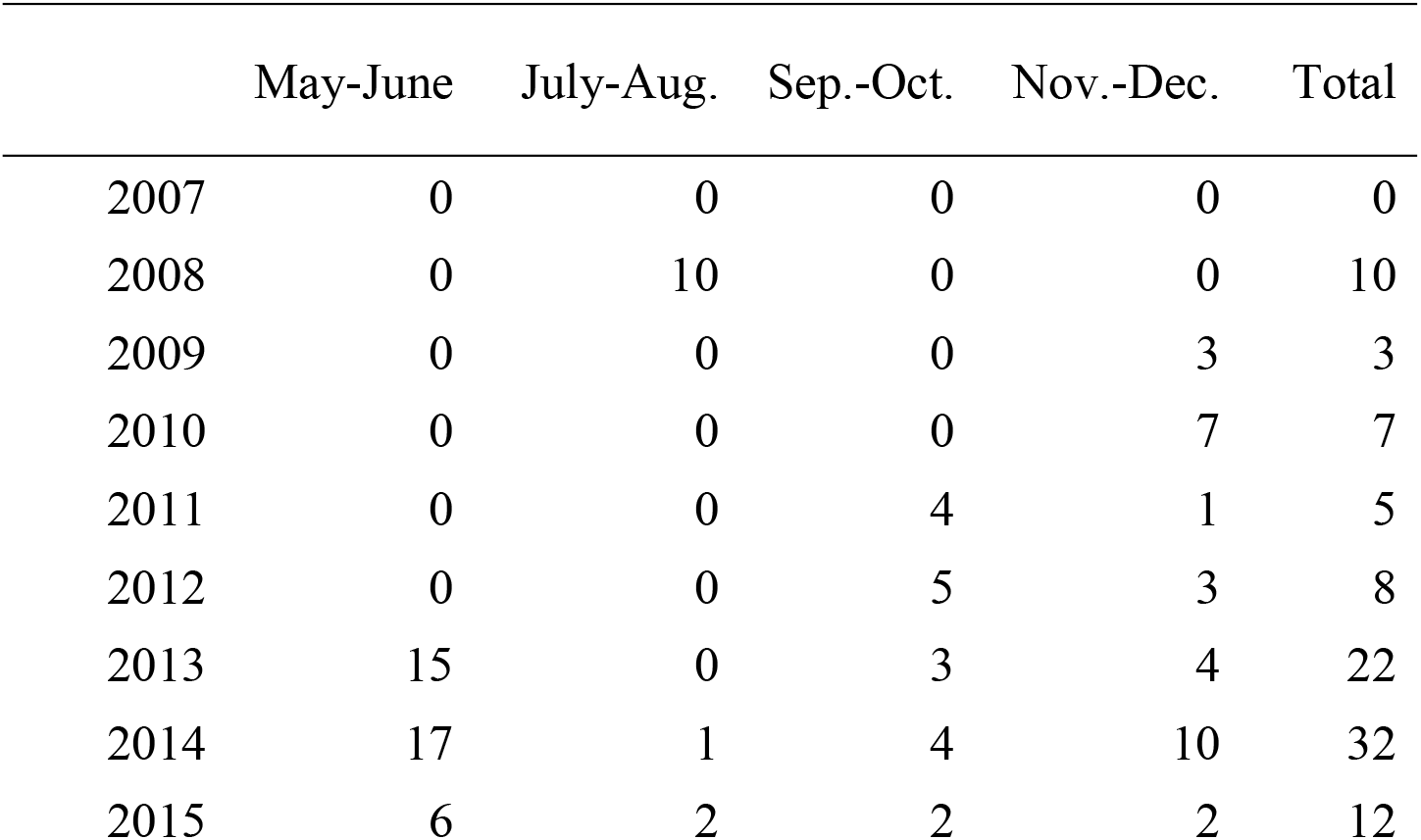
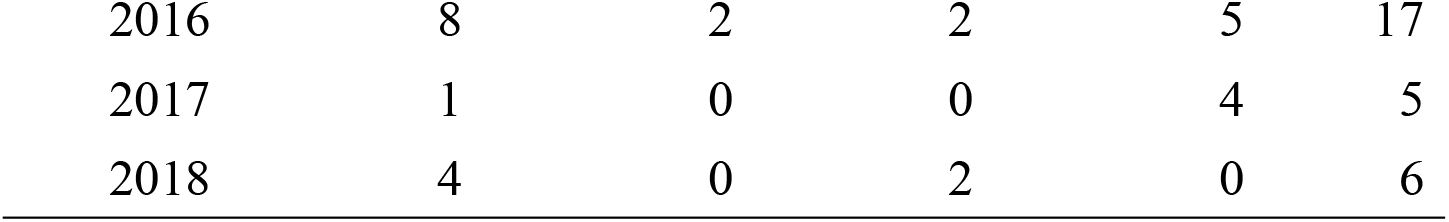
The numbers of deer hunted by nuisance control.

### Road count

We selected three route sectors (A: 4.7 km, B: 3.3 km, E: 0.7 km; Fig. 1) to record the numbers of deer sighted. The investigators of this study were mainly researchers and technical staff employed by the Forest Research Station, including non-specialists in deer. They recorded the date, weather, sector name, and time when they began driving through each sector, whenever they drove through a whole sector by vehicle during the period from May 1, 2007 to December 31, 2018. Then, they recorded the number of deer in each sector. If they found no deer, they recorded the number as zero. The details of the survey are described in a previous study [19].

We excluded records that lacked information about the number of deer sighted, sector name, year, and date. We also excluded records from January to April because few records were available from these periods due to snow accumulation and driving speed was different from other seasons. Furthermore, data within 15 minutes before and after were excluded from later analysis because data independence could not be guaranteed. After this data cleaning, we used 8,616 records for later analysis.

### Block count

Block counts were conducted in two sites (north: 86.9 ha, south: 111.7 ha) of the Ashiu Forest Research Station in December, from 2001 to 2018 except for 2017 (Fig 1). The sites were divided into 14 and 19 blocks (5 – 7 ha per block depending on the terrain), respectively. Each block was thoroughly surveyed by an observer walking in a zig-zag motion along the terrain in order to guarantee good visibility. When an observer spotted deer, they informed the observers of adjacent blocks using transceivers to avoid duplicate counting. Occasionally, we did not survey some blocks due to sudden snowfall and lack of observers; however, the total surveyed area was 181.27 ha in most years. Because it was a missing value only in 2017 and values did not change so much in the previous (2016) and next year (2018), we used the mean of 2016 and 2018 as the value of 2017 in the model described later.

### Number of deer carcasses at spring thaw

We counted the number of deer carcasses found in forest during the thawing period from April to early July, for the years 2005 – 2016. We needed to find deer carcasses emerging from the snow before animals preyed on them. We covered a 1 – 21 km distance per survey and repeated the procedure for 10 – 18 times per year to look for deer carcasses across the forest (Table 1 and S1 Fig.). In addition to looking for dead deer, we also relied on our sense of smell and detected carcasses based on the odor they emitted.

### State–space model

We analyzed field observation data of relative abundance indices of deer with state– space models, based on a hierarchical Bayesian framework [10, 25, 26]. The state–space model divided the observation data into a system model, representing “true” but unknown population size, and an observation model that accounts for error in counts caused by imperfect detection [25, 26]. The state–space models allowed us to permit potential errors in the count data. In most past studies in deer population dynamics, the analysis was performed on a yearly basis. However, we set the time interval to 2 months, excluding the period from January to April (*t* = 1 in May and June 2007, *t* = 2 in July and August 2007, *t* = 3 in September and October 2007, *t* = 4 in November and December 2007, *t* = 5 in May and June 2008, etc.). This was because we were able to use the road count data from all year round except from January to April (when the forest was covered by snow). We wanted to know the seasonal in addition to the annual fluctuation.

### System models

Deer abundance at time *t* (*N*_*t*_) in the forest depended on deer abundance at time *t* −1 (*N*_*t*-1_); the number varied with the effect of population growth (*r*_*t*_) including birth, natural mortality, immigration, migration at time *t* (*r*_*t*_ did not include the effects of hunting and mortality due to snowfall), and the effect of hunting at time *t* (hunting rate: *h*_*t*_). It can be expressed as follows:

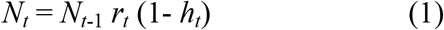

During the season when the forest was covered by snow (January to April), the deer sometimes got stuck or starved, due to lack of food as a result of the heavy snow. We defined the mortality rate during the seasons when the forest was covered by snow, just before the time *t*, as *d*_*t*_. Then, *N*_*t*_ (*t* = 5, 9, 13, ⋯45) can be expressed as follows:

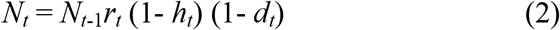

If we calculate the logarithm of the two aforementioned equations, then the process follows a linear structure. Then, equations (1) and (2) can be re-written as follows:

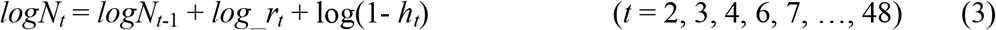

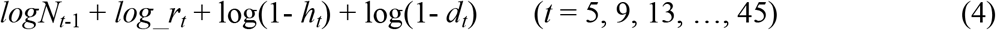

We introduced stochasticity into the deer population dynamics. Then, equation (3) can be expressed as follows:

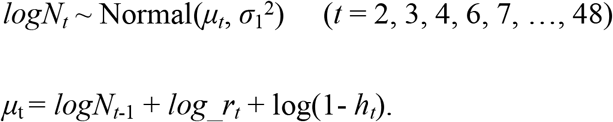

Equation (4) can be expressed as follows:

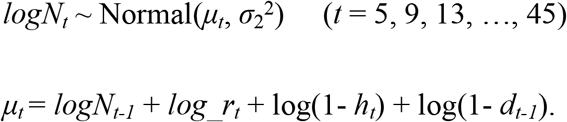

For the time interval we skipped four months every eight months, because we did not use the data collected from road count surveys by vehicles from the winter season (January to April). Thus, we defined different standard deviations of posterior distribution for deer abundance in the logarithmic scale (*σ*_1_ and *σ*_2_).

The prior probability distribution of the log of deer abundance in the first year (*logN*_1_) was determined as follows:

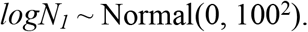

The population growth rate (logarithmic scale) at *t* was modeled as follows:

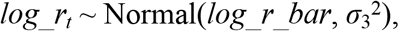

where *log_r_bar* is the mean population growth rate during the study period (logarithmic scale). We did not include a density-dependence parameter. Because the forest was not a closed ecosystem, we considered that the density-dependent decline in the birth rate of deer would rarely occur.

The hunting rate (logarithmic scale) at *t* was modeled as follows:

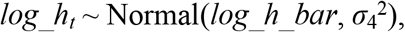

where *log_h_bar* is the mean hunting rate during the study period (logarithmic scale) and *σ*_4_ is the standard deviation of posterior distribution of the hunting rate in the logarithmic scale.

The prior probability distribution of *log_h_bar* was determined as follows:

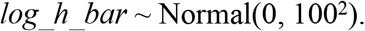

The mortality rate during seasons when the forest was covered by snow (logarithmic scale) at *t* (*log_d*) was modeled as follows:

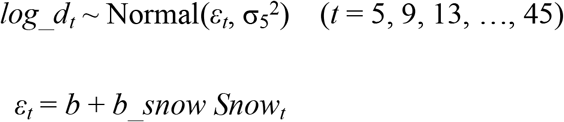

Where *ε*_*t*_ is the mean mortality during the winter with severe snowfall in time *t* and *σ*_5_ is the standard deviation of the posterior distribution of mortality during the winter with snowfall, in the logarithmic scale. Because the mortality may increase in severe snowfall conditions, it was assumed to increase linearly with the number of days with a snow depth of > 50 cm (*Snow*) before time *t* (logarithmic scale). To consider the different effects of snowfall, we also used the maximum snow depth instead of the number of days with snow depth of > 50 cm (S2 Table). The *b* and *b_snow* were the intercept and coefficient, respectively. The prior probability distributions of *b* and *b_snow* were as follows:

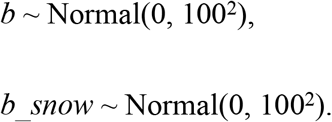

We assigned weakly informative priors for scale parameters, σ_1_ to σ_5_ as Cauchy(0, 10).

### Observation models

We modeled the number of deer seen in road count surveys (C_*tm*_) as follows:

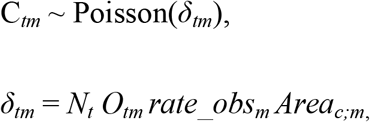

where *C*_*tm*_ is the mean number of deer seen in time *t* in route *m* (*m* = a, b, e), *rate_obs*_*m*_ is the observation rate per survey that converts *N*_*t*_ to *δ*_*tm*_, *O*_*tm*_ is the number of survey occasions conducted over two months for each route, and *Area*_*c;m*_ is the ratio of the study area in each drive count route (we assumed the census width to be 15m) per that of forest (a: 0.153%, b:0.107%, c: 0.023%). We assumed *C*_*tm*_ followed Poisson distribution, because the number of deer seen is a discrete, non-negative value. The prior probability distribution of *rate_obs_m_* were as follows:

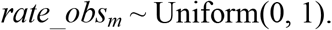

We modeled the number of deer seen by block count (*B*_*t*_) as follows:

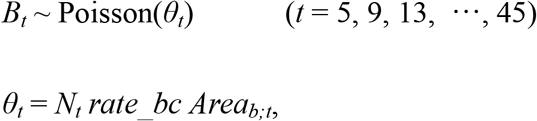

where *θ*_*t*_ is the mean number of deer seen by block count in time *t*, *rate_bc* is the observation rate per unit area that converts *N*_*t*_ to θ_*t*_, and *Area*_*b;t*_ is the survey area of the block count in time *t*. We assumed *B*_*t*_ followed a Poisson distribution, because block counts are discrete, non-negative values. The prior probability distribution of *rate_bc* was as follows:

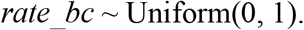

We modeled the number of deer hunted by nuisance control (*H*_*t*_) as follows:

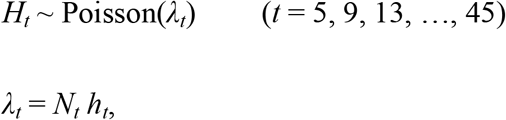

where *λ*_*t*_ is the mean number of deer hunted in time *t.* We assumed *H*_*t*_ followed a Poisson distribution, because hunted numbers are discrete, non-negative values. We modeled the number of deer carcasses found after thawing (*D*_*t*_) as follows:

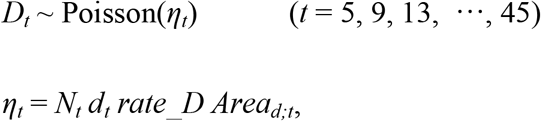

where *η*_*t*_ is the number of dead deer found after thawing in time *t*, *rate_D* is the detection rate per unit area that converts *N*_*t*_ *d*_*t*_ to *η*_*t*_, and *Area*_*d;t*_ is the survey area of dead deer surveyed after thawing in time *t*. We assumed *D*_*t*_ followed a Poisson distribution, because numbers of deer carcasses are discrete, non-negative values.

The parameter estimation was performed by the Markov Chain Monte Carlo (MCMC; [28]) calculation using RStan 2.18.2 [29]. We ran three parallel MCMC chains and retained 30,000 iterations after an initial burn-in of 15,000 iterations. We thinned sampled values to 1.0%. Convergence of MCMC sampling was judged by the criterion that the potential scale reduction factor on split chains, 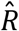 was smaller than 1.1 [30] and by a check of the MCMC trace.

## Results

The 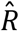 values of our estimated parameters were all under 1.1. The estimated deer abundance had two peaks (September–October 2007 and September–October 2010, Fig 2.) during the 12-year period. From 2011 to 2018, the estimated deer abundance was stable compared to the other periods. In most years, the models commonly indicated seasonal patterns of increase (spring to summer) and decrease (autumn to winter) in deer abundance. The mean of observation rates in route E was higher than that in routes A and B (Table 3). The mean of *σ*_1_ was lower than that of *σ*_2_. The 95% credible interval (CI) of *log_h_bar* was −6.31 to −4.44. On the other hand, the 95% CI of *log_r_bar* and *b_snow* included 0. Even when maximum snow depth was used instead of the number of days with snow depth of > 50 cm, the 95% CI of *b_snow* included 0.

**Fig 2.**
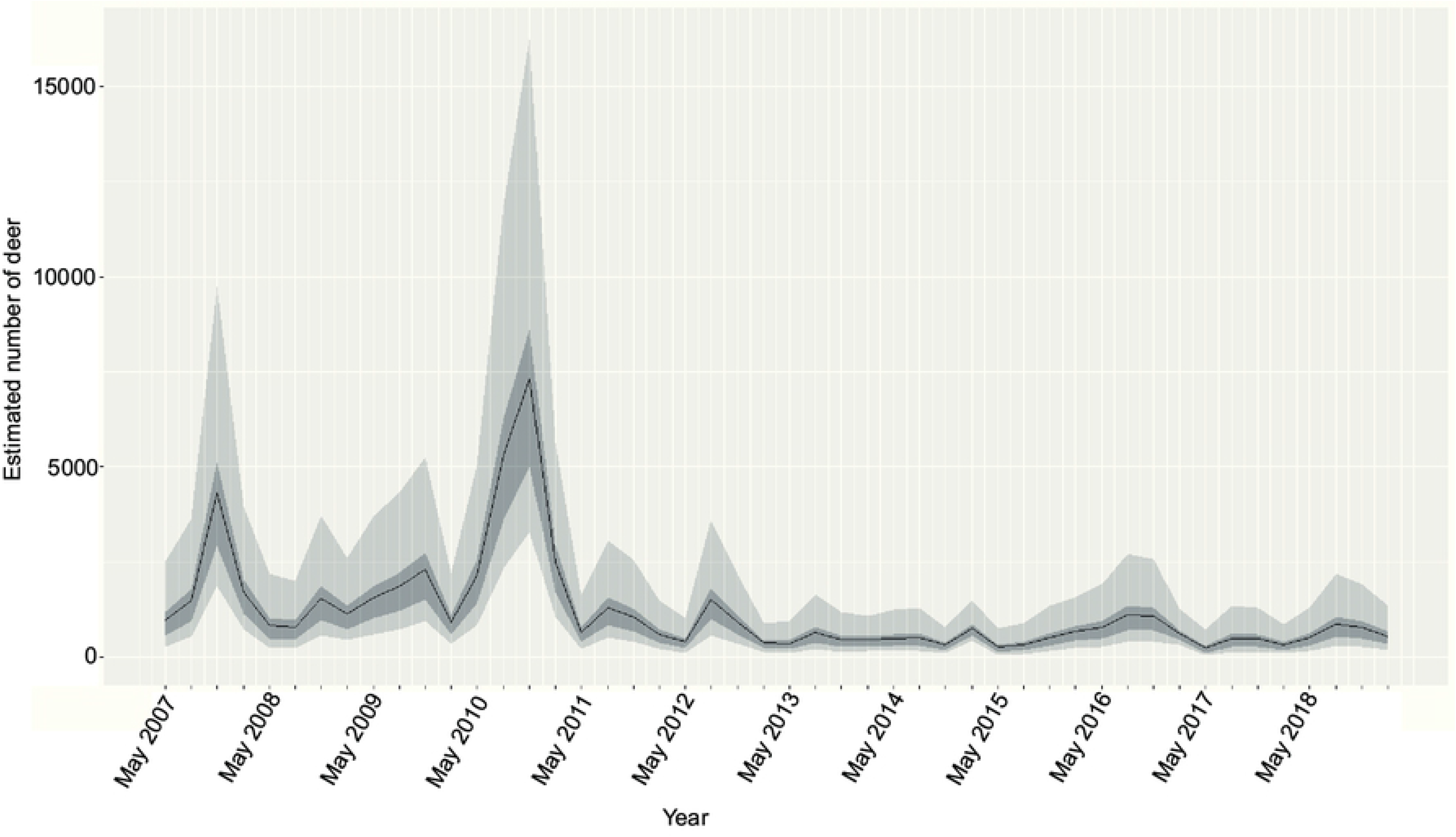
Estimated deer abundances that were obtained from the state–space model from 2008 to 2018. The black line denotes the mean of estimated deer abundance. The 50% and 95% credible intervals are denoted the dark and light gray, respectively.

**Table 3.**
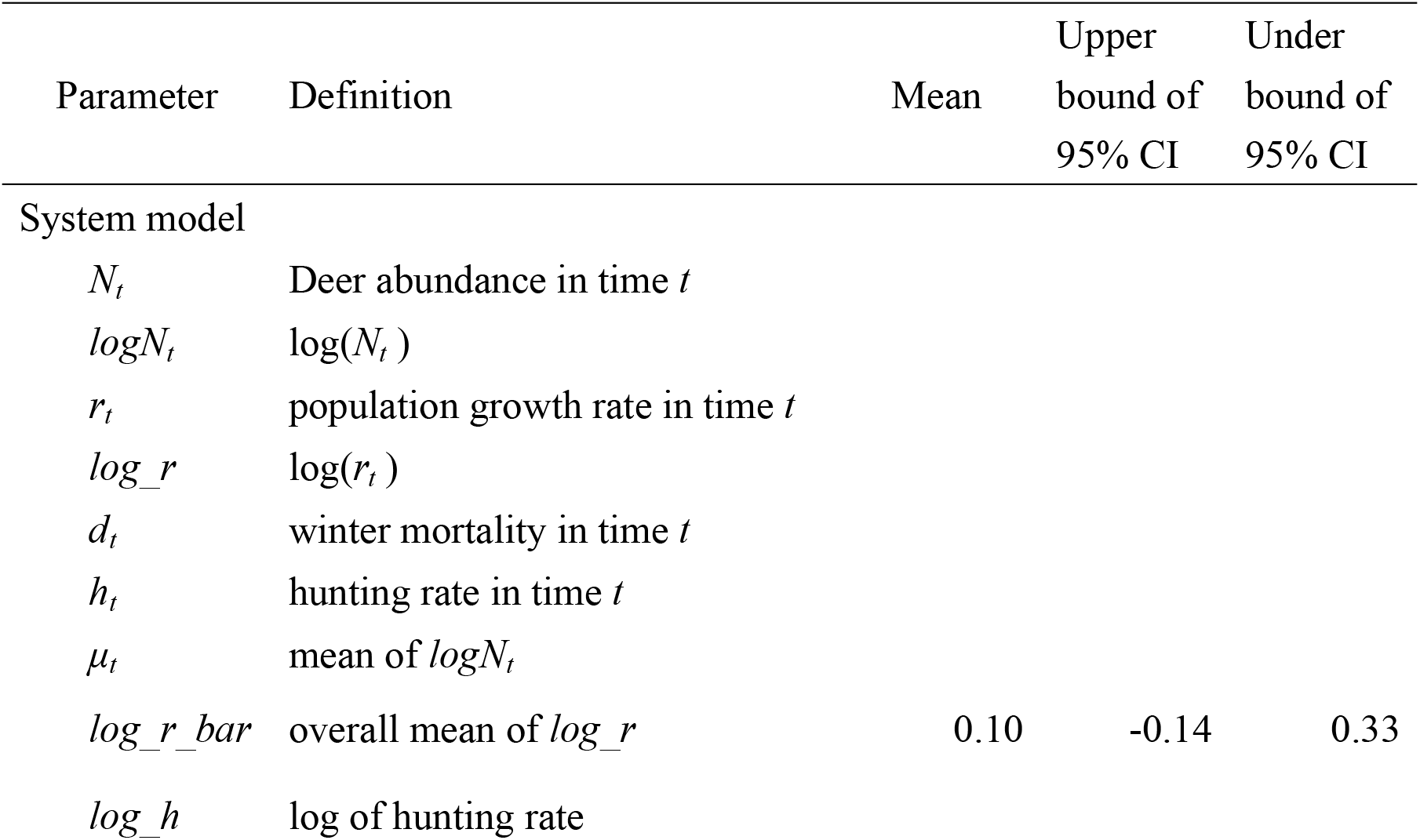
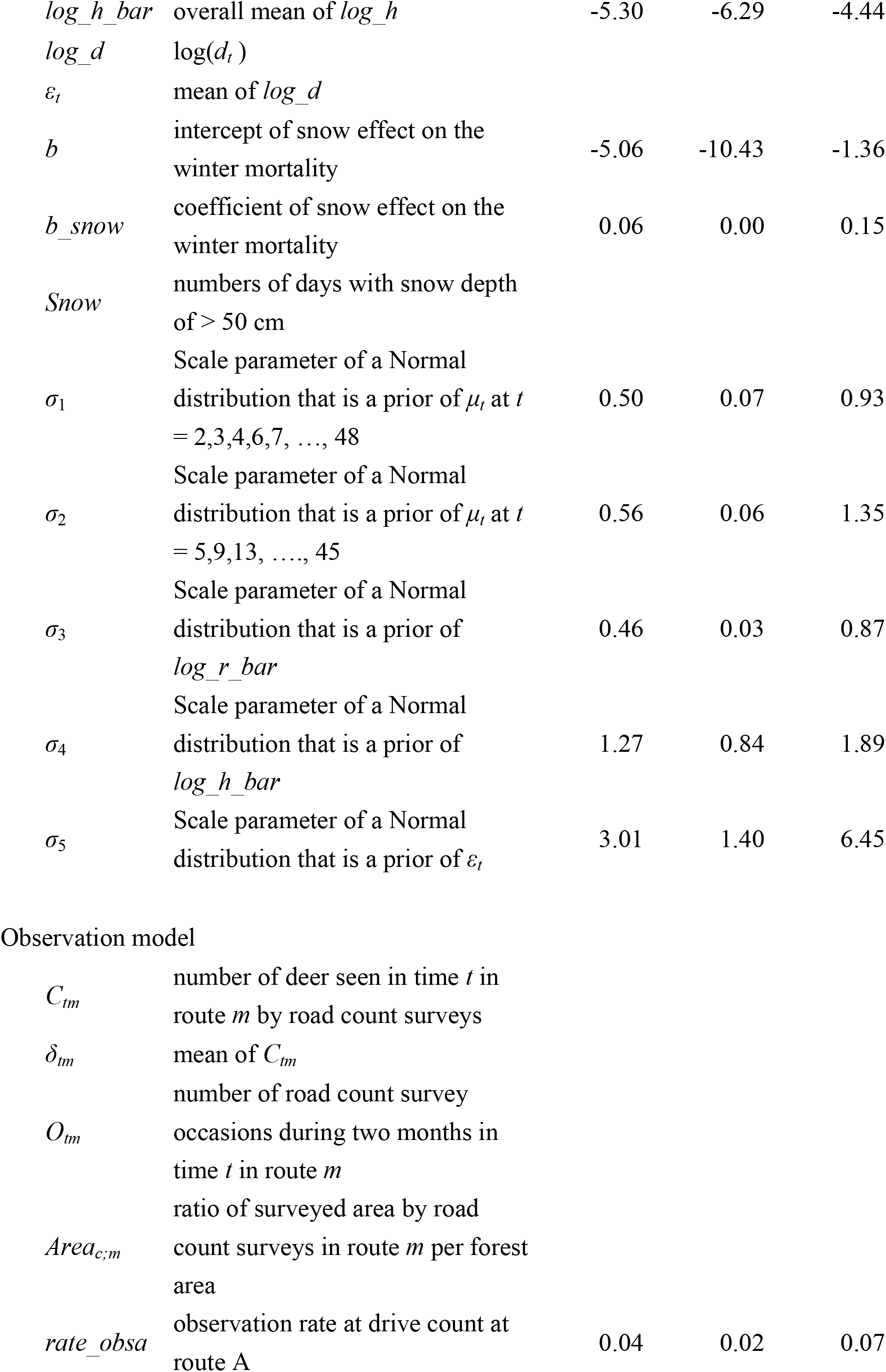
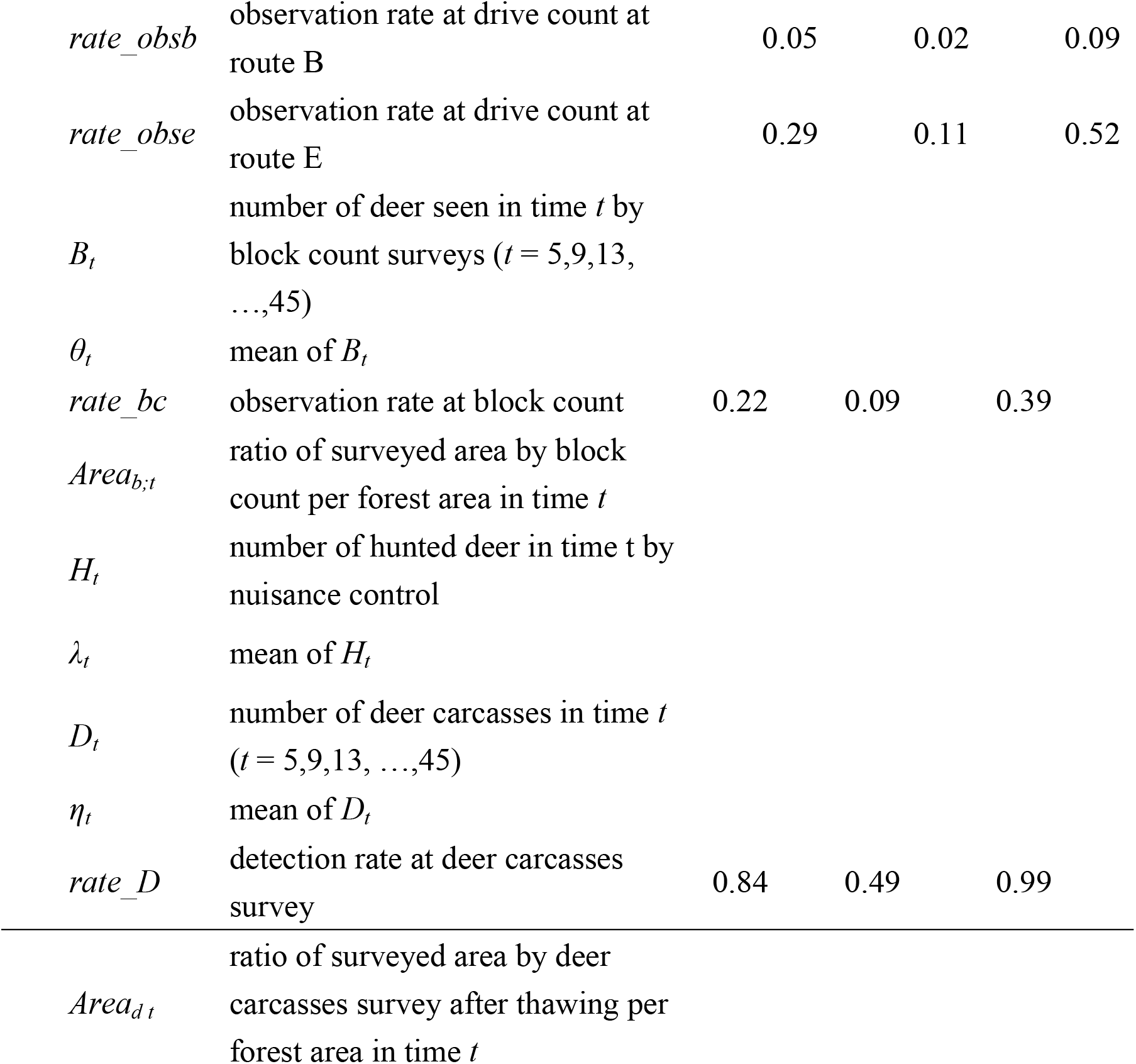
Data and parameters designed to estimate deer abundances from multiple abundance indices and posterior summaries of coefficients from the model.

| Parameter | Definition | Mean | Upper bound of 95% CI | Under bound of 95% CI |
| --- | --- | --- | --- | --- |
| System model |  |  |  |  |
| $N_t$ | Deer abundance in time $t$ | | | |
| $\log N_t$ | $\log(N_t)$ | | | |
| $r_t$ | population growth rate in time $t$ | | | |
| $\log\_r$ | $\log(r_t)$ | | | |
| $d_t$ | winter mortality in time $t$ | | | |
| $h_t$ | hunting rate in time $t$ | | | |
| $\mu_t$ | mean of $\log N_t$ | | | |
| $\log\_r\_bar$ | overall mean of $\log\_r$ | 0.10 | -0.14 | 0.33 |
| $\log\_h$ | log of hunting rate | | | |
| $\log\_h\_bar$ | overall mean of $\log\_h$ | -5.30 | -6.29 | -4.44 |
| $\log\_d$ | $\log(d_t)$ | | | |
| $\varepsilon_t$ | mean of $\log\_d$ | | | |
| $b$ | intercept of snow effect on the winter mortality | -5.06 | -10.43 | -1.36 |
| $b\_snow$ | coefficient of snow effect on the winter mortality | 0.06 | 0.00 | 0.15 |
| $Snow$ | numbers of days with snow depth of > 50 cm | | | |
| $\sigma_1$ | Scale parameter of a Normal distribution that is a prior of $\mu_t$ at $t = 2, 3, 4, 6, 7, \dots, 48$ | 0.50 | 0.07 | 0.93 |
| $\sigma_2$ | Scale parameter of a Normal distribution that is a prior of $\mu_t$ at $t = 5, 9, 13, \dots, 45$ | 0.56 | 0.06 | 1.35 |
| $\sigma_3$ | Scale parameter of a Normal distribution that is a prior of $\log\_r\_bar$ | 0.46 | 0.03 | 0.87 |
| $\sigma_4$ | Scale parameter of a Normal distribution that is a prior of $\log\_h\_bar$ | 1.27 | 0.84 | 1.89 |
| $\sigma_5$ | Scale parameter of a Normal distribution that is a prior of $\varepsilon_t$ | 3.01 | 1.40 | 6.45 |
| Observation model |  |  |  |  |
| $C_{tm}$ | number of deer seen in time $t$ in route $m$ by road count surveys | | | |
| $\delta_{tm}$ | mean of $C_{tm}$ | | | |
| $O_{tm}$ | number of road count survey occasions during two months in time $t$ in route $m$ | | | |
| $Area_{c,m}$ | ratio of surveyed area by road count surveys in route $m$ per forest area | | | |
| $rate\_obsa$ | observation rate at drive count at route A | 0.04 | 0.02 | 0.07 |
| $rate\_obsb$ | observation rate at drive count at route B | 0.05 | 0.02 | 0.09 |
| $rate\_obse$ | observation rate at drive count at route E | 0.29 | 0.11 | 0.52 |
| $B_t$ | number of deer seen in time $t$ by block count surveys ($t = 5, 9, 13, \dots, 45$) | | | |
| $\theta_t$ | mean of $B_t$ | | | |
| $rate\_bc$ | observation rate at block count | 0.22 | 0.09 | 0.39 |
| $Area_{b,t}$ | ratio of surveyed area by block count per forest area in time $t$ | | | |
| $H_t$ | number of hunted deer in time $t$ by nuisance control | | | |
| $\lambda_t$ | mean of $H_t$ | | | |
| $D_t$ | number of deer carcasses in time $t$ ($t = 5, 9, 13, \dots, 45$) | | | |
| $\eta_t$ | mean of $D_t$ | | | |
| $rate\_D$ | detection rate at deer carcasses survey | 0.84 | 0.49 | 0.99 |
| $Area_{dt}$ | ratio of surveyed area by deer carcasses survey after thawing per forest area in time $t$ | | | |

## Discussion

Using a Bayesian state–space model, we were able to estimate annual and seasonal fluctuations of deer abundance with data collected from block count surveys, road count surveys by vehicles, mortality surveys during the winter, and nuisance control. Individual indices have their own biases against certain environmental factors, but the effect of these biases can be mitigated by using multiple indicators [10, 31, 32]. The estimated standard deviation of posterior distribution for deer abundance in the logarithmic scale was higher in May and June, immediately after winter, than it was in other seasons (*σ*_2_ > σ_1_, Table 3). In our study, the time interval we used skipped four months every eight months, because we did not use the data of road count surveys by vehicles from the winter season (January to April). Thus, deer abundance after winter could vary more than during other time intervals, leading to higher estimated variance. By using a Bayesian state–space model, we detected this kind of heterogeneity in variance.

The mean deer abundance in the forest in 2014 was estimated to be 490–766 individuals per 4,612 ha (Fig 2), namely, 10.6–16.6 individuals per km^2^. The 95% CI of this estimation corresponds to 4.0–18.9 individuals per km^2^. The Japanese Ministry of the Environment published a density distribution map of deer in Japan in 2014 complied by using fecal pellet group counts and estimated deer abundance by using a hierarchical Bayesian model that took into account the number of hunted deer [33]. According to the report, deer density in the mesh (25 km^2^/mesh) our study site belongs to was estimated as 7–10 individuals per km^2^ in 2014. Thus, our estimate was similar to that of the Ministry. Although it is unclear which estimate is closer to the true value, it is important to focus on patterns of deer population dynamics rather than the values themselves. We found two peaks of deer abundance during the 12-year study period (September– October, 2007 and September–October, 2010, Fig 2). The estimated deer abundance at the autumn of 2010, in particular, was the highest (159.7 individuals per km^2^). The peak in 2010 could be considered an outbreak; this is also reported in other populations and deer species [17, 34]. In 2011, the estimated deer abundance decreased drastically, and has remained at a low level since then. By 2003, most shrubs, herbs, and dwarf bamboo in the forest had already been overgrazed [23, 35, 36]. Therefore, the 2010 irruption and the 2011 decrease could not be due to food shortage of the understory vegetation. When the understory vegetation was poor, deer may have been depending strongly on nuts from canopy and sub-canopy species as food sources during the autumn. In the autumn of 2009, nut production was synchronously very high in three dominant masting Fagaceae species (*Fagus crenata*, *Quercus crispula* and *Quercus serrata*) in the Hyogo prefecture that lies next to the prefecture the study forest belongs to; then, nut production was synchronously very low in the autumn of 2010 [37]. Although we did not collect any masting data from our study site, a nut shortage may have affected the drastic deer population decrease of 2011. From 2011, the aforementioned three tree species did not produce nuts synchronously. This asynchronous nut production might have led to low deer population stability starting from 2011 onwards.

In this study, the seasonal fluctuation of deer abundance tended to be similar to past results obtained from road count surveys by vehicles [38]. Deer abundance mainly increased from spring to autumn and decreased from autumn to winter (Fig 2). Though some deer exist in forests even during the winter [37], they migrate seasonally to avoid snow accumulation in heavy snow-covered areas [39, 40]. Therefore, the seasonal variation we detected may be due to the seasonal migration pattern in addition to the population recruitment through fawn births in early summer. The potential browsing pressure increase in the plant community during the summer may have negative effects on herbaceous plants, especially the one that grow in the summer and flowered in the autumn. In this area, as the plants that flower after midsummer are herbaceous and are more severely browsed compared to trees [41], the fitness of pollinators working from summer to autumn may critically decrease due to a shortage in their flower resources. The 95% CI of *log_h_bar* ranged from −6.29 to −5.30, though the 95% CI of *log_r_bar* included 0 (Table 3). These results suggest that nuisance control could be useful in decreasing deer populations and are similar to past results [10]. On the other hand, a previous study [10] pointed out the difficulties of increasing hunting pressures because Japanese hunters were getting older. To establish an effective deer abundance management program under this circumstance, the development of simple and inexpensive capture methods is urgent.

Late snowfall substantially affects the mortality of *C. nippon* [12]. In *Cervus elaphus* in Norway, winter harshness affects first-year survival but not the survival of adults [42]. In this study, the 95% CI of *b_snow* included 0. This suggests that snowfall may have slightly affected deer mortality during the winter in the present study. This is similar to results obtained from studying the alpine ungulate *Rupicapra rupicapra* [15], though their population dynamics are largely affected by summer temperature. At first glance, our results seem to suggest that the mortality rate during the winter will not change even if snowfall decreases due to global warming. However, as we mentioned earlier, deer inhabiting regions with heavy snowfall, migrate to safe areas during the winter and go back to their initial habitats after snowmelt. Thus, snowfall decrease due to global warming may decelerate the winter migration of deer and, subsequently, deer that remain on-site may intensively forage evergreen perennial plants during the winter season.

In route E, the observation rate was higher than that in routes A and B (Table 3). In this study, we did not consider the spatial pattern of deer. While route A is close to a village, route E is remote and located deep in montane forest. Therefore, human activity may have affected the observation rate. The topographic pattern could have affected route visibility, though we uniform ranges of observation 15 m width in all routes. Landscape characteristics such as evergreen forests and artificial grasslands affect deer abundance in local areas [10]. As shown in Fig 1, this study site consists of steep slopes and deep valleys. The differences in observation rates among routes may also be due to the differences in landscape characteristics in a local scale in the forest.

In conclusion, we clarified the population dynamics of deer not only annually but also seasonally. Snowfall accumulations did not affect population dynamics of deer in this study irrespective of higher mortality of deer during the winter [9, 12]. However, we need to pay attention to the effect the winter migration of deer has on plant communities because many deer migrated to another area during the winter and came back before the summer. Although we could not grasp the population dynamics during the snow accumulation season, in warmer winters, more deer may remain in the forest. Thus, a warmer winter may lead to degradation of evergreen perennial plant communities during the winter and early spring. Additional investigation on evergreen perennial plants could help examine the effect of deer browsing during the winter. In contrast to snowfall accumulations, nuisance control had an effect on the population dynamics of deer. Even in wildlife protection areas and national parks where hunting is regulated, nuisance control could be effective in buffering the effects of excessive deer browsing on forest ecosystems as well as plant communities, under the absence of potent predators.

## Acknowledgments

The authors would like to thank all the members who participated in this monitoring study and the staff of the Ashiu Forest Station for their support.

## Supporting information

**S1 Fig. Survey route where we counted the number of deer carcasses found in forest during the thawing period from April to early July, for the years 2017 – 2018.** The parts surrounded by solid lines denote the area of Ashiu Forest Station and the area surrounding. The red lines denote the survey route of the year. The routes are slightly different from year to year.

**S1 Table. Numbers of deer observed (A, B, E) and numbers of survey (obA, obB, obC) by road count surveys at each sector, respectively.**

**S2 Table. Numbers of deer observed and area by block count surveys.**

**S3 Table 3. Data and parameters designed to estimate deer abundances from multiple abundance indices and posterior summaries of coefficients from the model which used MaxS instead of SD50 in Table 1 as snow effect.**

**S1 Appendix. R-Code for fitting the Bayesian state-space model.** Simulation code using a Bayesian state–space model.

**S1 Dataset.**

## References

1. Gill RMA. A review of damage by mammals in north temperate forests: 1. Deer. Forestry. 1992;65:145–69.

2. Takatsuki S. Effects of sika deer on vegetation in Japan: A review. Biol Conserv. 2009;142:1922–1929.

3. Akashi N, Nakashizuka T. Effects of bark-stripping by Sika deer (Cervus nippon) on population dynamics of a mixed forest in Japan. For Ecol Manage. 1999;113:75–82.

4. Fujiki D, Kishimoto Y, Sakata H. Assessing decline in physical structure of deciduous hardwood forest stands under sika deer grazing using shrub-layer vegetation cover. J For Res. 2010;15:140–144.

5. Fuller RJ, Gill RMA. Ecological impacts of increasing numbers of deer in British woodland. Forestry. 2001;74:193–199.

6. Filazzola A, Tanentzap AJ, Bazely DR. Estimating the impacts of browsers on forest understories using a modified index of community composition. For Ecol Manage. 2014;313:10–16.

7. Hobbs NT, Swift DM. Estimates of habitat carrying-capacity incorporating explicit nutritional constraints. J Wildl Manage. 1985;49:814–822.

8. Kaji K, Koizumi T, Ohtaishi N. Effects of resource limitation on the physical and reproductive condition of sika deer on Nakanoshima island, Hokkaido. Acta Theriol. 1988;33:187–208.

9. Ueno M, Iijima H, Takeshita K, Takahashi H, Yoshida T, Uehara H, et al. Robustness of adult female survival maintains a high-density sika deer (Cervus nippon) population following the initial irruption. Wildl Res. 2018;45:143–154.

10. Iijima H, Nagaike T, Honda T. Estimation of deer population dynamics using a bayesian state–space model with multiple abundance indices. J Wildl Manage. 2013;77:1038–1047.

11. Hagen R, Haydn A, Suchant R. Estimating red deer (*Cervus elaphus*) population size in the Southern Black Forest: the role of hunting in population control. Eur J Wildl Res. 2018 Aug;64:doi: 10.1007/s10344-018-1204-z.

12. Takatsuki S, Suzuki K, Suzuki I. A mass-mortality of Sika deer on Kinkazan Island, northern Japan. Ecol Res. 1994;9:215–223.

13. Kawase H, Sasaki H, Murata A, Nosaka M, Ishizaki NN. Future Changes in Winter Precipitation around Japan Projected by Ensemble Experiments Using NHRCM. J Meteorol Soc Jpn. 2015;93:571–580.

14. Bonardi A, Corlatti L, Bragalanti N, Pedrotti L. The role of weather and density dependence on population dynamics of Alpine-dwelling red deer. Integr Zool. 2017;12:61–76.

15. Ciach M, Pęksa Ł. Impact of climate on the population dynamics of an alpine ungulate: a long-term study of the Tatra chamois Rupicapra rupicapra tatrica. Int J Biometeorol. 2018;62:2173–2182.

16. Takeshita K, Tanikawa K, Kaji K. Applicability of a Bayesian state–space model for evaluating the effects of localized culling on subsequent density changes: sika deer as a case study. Eur J Wildl Res. 2017 Aug;63:doi: 10.1007/s10344-017-1128-z.

17. Takeshita K, Ueno M, Takahashi H, Ikeda T, Mitsuya R, Yoshida T, et al. Demographic analysis of the irruptive dynamics of an introduced sika deer population. Ecosphere. 2018 Sep 13;9(9):e02398. doi: 10.1002/ecs2.2398.

18. Pellerin M, Bessiere A, Maillard D, Capron G, Gaillard JM, Michallet J, et al. Saving time and money by using diurnal vehicle counts to monitor roe deer abundance. Wildl Biol. 2017 Jan 1;(4):doi: 10.2981/wlb.00274.

19. Mizuki I, Sakaguchi S, Fukushima K, Sakai M, Takayanagi A, Fujiki D, et al. Among-year variation in deer population density index estimated from road count surveys. J For Res. 2013;18:491–497.

20. Valente AM, Marques TA, Fonseca C, Torres RT. A new insight for monitoring ungulates: density surface modelling of roe deer in a Mediterranean habitat. Eur J Wildl Res. 2016;62:577–587.

21. Trenkel VM, Elston DA, Buckland ST. Fitting population dynamics models to count and cull data using sequential importance sampling. J Am Stat Assoc. 2000;95:363–374.

22. ASHIU FOREST RESEARCH STATION [Internet]. Field Science Education and Research Center, Kyoto University. 2019 [cited 25 Oct 2019]. Available from: http://fserc.kyoto-u.ac.jp/wp/blog/archives/945.

23. Kato M, Okuyama Y. Changes in the biodiversity of a deciduous forest ecosystem caused by an increase in the Sika deer population at Ashiu, Japan. Contr Biol Lab Kyoto Univ. 2004;29:437–448.

24. Sakaguchi S, Fujiki D, Inoue M, Yamasaki M, Fukushima K, Takayanagi A. Plant species preference of sika deer in cool-temperate mixed conifer-broadleaf forest of the Sea of Japan side of Central Japan. For Res Kyoto. 2012;78:71–80.

25. Clark JS, Bjørnstad ON. Population time series: Process variability, observation errors, missing values, lags, and hidden states. Ecology. 2004;85:3140–3150.

26. Marshal JP, Rankin C, Nel HP, Parrini F. Drivers of population dynamics in sable antelope: forage, habitat or competition? Eur J Wildl Res. 2016;62:549–56.

27. Kaji K, Okada H, Yamanaka M, Matsuda H, Yabe T. Irruption of a colonizing sika deer population. J Wildl Manage. 2004;68:889–899.

28. Calder C, Lavine M, Muller P, Clark JS. Incorporating multiple sources of stochasticity into dynamic population models. Ecology. 2003;84:1395–1402.

29. Stan Development Team. RStan. Version 2.18.2 [software]. 2019 Oct 25 [cited 2019 Oct 25]. Available from : http://mc-stan.org.

30. Gelman A, Carlin JB, Stern HS, Dunson DB, Vehtari A, Rubin DB. Bayesian Data Analysis. 3rd ed. Florida: Chapman & Hall/CRC; 2013.

31. Uno H, Kaji K, Saitoh T, Matsuda H, Hirakawa H, Yamamura K, et al. Evaluation of relative density indices for sika deer in eastern Hokkaido, Japan. Ecol Res. 2006;21:624–632.

32. Morellet N, Gaillard JM, Hewison AJM, Ballon P, Boscardin Y, Duncan P, et al. Indicators of ecological change: new tools for managing populations of large herbivores. J Appl Ecol. 2007;44:634–643.

33. Ministry of the Environment. Government of Japan [Internet]. Env.go.jp. 2019 [cited 2019 Oct 25]. Available from: https://www.env.go.jp/nature/choju/conf/conf_wp/conf02-h29/ref01.pdf.

34. Coulson T, Guinness F, Pemberton J, Clutton-Brock T. The demographic consequences of releasing a population of red deer from culling. Ecology. 2004;85:411–422.

35. Fukuda A, Takayanagi A. Influence of snow cover on browsing of Cephalotaxus harringtonia var. nana by *Cervus nippon* Temminck in heavy snow region in central Japan. For Res Kyoto. 2008;77:5–11.

36. Tanaka Y, Takatsuki S, Takayanagi A. Decline of Sasa palmata community by grazing of Sika deer (*Cervus nippon*) at Ashiu Research Forest Station. For Res Kyoto. 2008;77:13–23.

37. Fujiki D. Can frequent occurrence of Asiatic black bears around residential areas be predicted by a model-based mast production in multiple Fagaceae species? J For Res. 2018;23:260–269.

38. Goda R, Mizuki I, Takayanagi A. Road census for sika deer monitoring at the Ashiu Forest Research Station of Kyoto University. For Res Kyoto. 2008;77:89–94.

39. Igota H, Sakuragi M, Uno H, Kaji K, Kaneko M, Akamatsu R, et al. Seasonal migration patterns of female sika deer in eastern Hokkaido, Japan. Ecol Res. 2004;19:169–178.

40. Takii A, Izumiyama S, Taguchi M. Partial migration and effects of climate on migratory movements of sika deer in Kirigamine Highland, central Japan. Mamm Study. 2012;37:331–340.

41. Kato M, Kakutani T, Inoue T, Itino T. Insect-flower relationship in the primary forest of Ashu, Kyoto: An overview of the flowering phenology and the seasonal pattern of insect visits. Contr Biol Lab Kyoto. 1990;27:309–375.

42. Loison A, Langvatn R. Short- and long-term effects of winter and spring weather on growth and survival of red deer in Norway. Oecologia. 1998;116:489–500

